# One scale does not fit all: invasive predator identity determines the impact on native prey

**DOI:** 10.64898/2026.06.26.734748

**Authors:** Albert Bonet Bigata, Chris Sutherland, Xavier Lambin

## Abstract

1. When eradication is unfeasible, invasive predator control should evaluate how removal affects ecological responses by native species. Assessments often use total invasive predator abundance to evaluate prey responses, yet intraspecific variation in diet and space use means that some subgroups cause disproportionate impacts. Identifying these ‘problem individuals’, and the spatial scales over which their impacts operate, can enable targeted spatially explicit removal to maximise impact reduction. However, despite individual-level information is often already collected during trapping operations it is seldom included when analysing predator impacts, potentially biasing the conservation outcomes expected under blanket removal.
2. We use a novel framework and two decades of invasive predator control data to estimate how individual variation in residency status influences the distance-dependent impacts of invasive American mink *Neogale vison* on water vole *Arvicola amphibius* occupancy across two prey surveys. We also develop a sub-model to predict mink residency status for individuals with missing age data.
3. The probability of capturing adult mink decreased with elevation and years of control, indicating that long-term control altered the resident population and demographic composition of mink around water vole sites.
4. Distance-dependent negative impacts of mink varied by residency status, becoming negligible at approximately 20 km from water vole sites for resident mink and 2 km for transient. The spatial scale of mink impacts was largest during the first vole survey when resident mink were more abundant, and declined rapidly for the second survey, when mink were less abundant and spatially clustered. Our results suggest that water voles have benefited mostly from reducing resident mink rather than the total population, especially in early control phases.
5. Managers can use our framework to develop spatially explicit and impact-based strategies, not restricted to invasive species control, to construct empirically informed management buffers around populations of conservation concern. Long-term efforts will change the landscape and invasive predator contexts, and thus we recommend iteratively updating and re-evaluating management outcome evaluations. We argue that incorporating individual heterogeneity improves our understanding of ecological mechanisms influencing management success but that the suitability of targeted strategies should be evaluated for target socioecological contexts.

## Introduction

Invasive non-native mammalian predators (‘invasive predators’, hereafter) are major contributors to global vertebrate defaunation (Doherty et al., 2016). Reducing the negative impact of invaders, often through lethal control, is a key objective of conservation programmes worldwide. Evidence-based invasive predator management requires information beyond invader removal, including the responses from native biota to removal efforts to assess and optimise the effectiveness of conservation actions (Doherty & Ritchie, 2017; W. J. Sutherland et al., 2004). Such ecological impacts of invasive species removal are seldomly reported in practice, limiting the ability to infer conservation gains from removal efforts (García-Díaz et al., 2021; Prior et al., 2018; but see Bird et al., 2024; Fox et al., 2025; Norbury et al., 2015).

Local ecological responses (e.g., changes in distribution, abundance, reproduction) of native species to removal depend on environmental features (e.g. habitat availability) and changes in invasive predator abundance (Fox et al., 2025; Nordström et al., 2002). Existing approaches for modelling abundance-impact relationships often use sympatric invasive predator abundance, overlooking the spatial distribution of predators beyond focal native prey sites. Additionally, they commonly aggregate all individuals into single metrics (Norbury et al., 2015; Yokomizo et al., 2009), assuming homogenous intraspecific predator impacts on native species.

Both decisions may be biasing outcome evaluations. Predation effects are inherently distance-dependent, with terrestrial predators operating at larger scales than their prey (Sutherland et al., 2012) Ignoring predators beyond prey sites but within the spatial range over which they influence prey can overestimate local predator impacts, thereby biasing predicted removal benefits and misinforming management outcomes. At the same time, the spatial scale of invasive predator space use, and thus the distance over which they influence native prey, is not uniform within invasive predators. Space use varies seasonally, with prey availability and with presence of apex predators (Garvey et al., 2022; Roshier & Carter, 2021; Salo et al., 2008). Individual foraging strategies (García-Díaz et al., 2021) and other traits such as sex (Wysong et al., 2020), reproductive status (Chinn et al., 2023; Dalal et al., 2021), and size can further influence predator space use, diet and consumption rates, leading to heterogeneous intraspecific impacts. For example, Moseby et al., (2015) demonstrated that larger male cats had disproportionately more frequent predation leading to the development of targeted size-based control strategies. Thus, assuming ‘one scale fits all’ when estimating distance-dependent invasive predator impacts, may overlook intraspecific heterogeneity and underestimate the *per capita* impacts of disproportionately damaging subgroups (‘problem individuals’ hereafter), which should be removed with higher priority.

Estimating how intraspecific variation in invasive predators space use translates into ecological impacts often relies on individual movement data (e.g., resource selection functions: Wysong et al., 2020; kernel density estimators: Maestresalas et al., 2023). This, typically, requires GPS or VHF telemetry, which is prohibitively expensive and suffers from small sample sizes (i.e., unique captured individuals) which limits the ability to make population-level inferences (Hebblewhite & Haydon, 2010). Non-invasive approaches, including environmental DNA (eDNA; Williams et al., 2018) and camera trapping (Garvey et al., 2022) have also been used but can be resource intensive or inaccurate, potentially diverting resources from control efforts (Bogich et al., 2008; Buxton et al., 2020).

One cost-effective alternative is to couple spatially explicit individual-level invasive predator data already collected in most trapping-based removal efforts with native prey monitoring. When collected concurrently in space and time, the distance between invasive predator captures and native prey sites can be used to estimate distance-dependent predation impacts (Mönkkönen et al., 2007). By treating predation as a landscape effect, spatially-explicit methods can be used to estimate how far in space negative invasive predator impacts span (e.g. (Chandler & Hepinstall-Cymerman, 2016; Gallo et al., 2018; Miguet et al., 2017). These estimates can then be used to design empirically informed spatially explicit management of both invasive predator and native prey species simultaneously. Despite their potential, such spatially explicit approaches have not been used in practice to estimate the spatial scale of negative invasive predators impacts nor identify ‘problem individuals’ for targeted removal.

In this study we aim to showcase the benefits of these spatially explicit methods by developing an analytical framework to quantify intraspecific heterogeneity in distance-dependent impacts of an invasive predator (American mink *Neogale vison*) on the distribution of a native prey, the water vole *Arvicola amphibius*.

American mink (‘mink’ hereafter) predation negatively impacts invertebrate and vertebrate species outside their native range (Bonesi & Palazon, 2007; Vada et al., 2023), particularly on ground and cliff-nesting birds and small mammals. A clear example is the contribution of mink to the >90% decline of water voles in the UK (Strachan et al., 2000).

Mink are semiaquatic generalist predators with pronounced sex, size and ontogenetic-specific variation in diet and home range sizes, with limited intrasexual overlap (Magnusdottir et al., 2014; Melero et al., 2008; Yamaguchi & Macdonald, 2003). Such intraspecific heterogeneity makes them a model species for testing how individual traits influence the scale of invasive predator impacts.

We hypothesize that the spatial scale of mink impacts on water vole populations will differ between adult mink holding breeding territories (‘resident’) and non-settled mink in dispersal or transient phases (‘transient’). Specifically, we expect that resident mink impacts will span over larger distances than transient mink due to cumulative predation on fragmented metapopulations linked by dispersal, such as water voles in the UK. Predation depletes source subpopulations that contribute to (re)colonization whilst increasing local and metapopulation-wide extinction probabilities (Holyoak & Lawler, 1996; Woodroffe et al., 1990). In contrast, while transient individuals may move large daily distances in search of a settlement area, we expect their impacts on local water vole populations to be spatially limited since they will spend less time predating within the same location. While this has not been studied in invasive mustelids, spatially clustered predation by resident predators and episodic, or spatially diffuse, by transient individuals has been shown in native predators such as Coyotes *Canis latrans* in North America (Chamberlain et al., 2021).

To test this, we use two decades of landscape scale mink capture data collected by non-professional volunteers as part of a large-scale control program (Lambin et al., 2019) in combination to two snapshot water vole surveys conducted at the end of the initial depletion phase and at the suppression stages of mink control respectively.

We demonstrate the value of linking spatially explicit predator impact heterogeneity to native prey responses using a novel framework for quantifying whether ecological outcomes of invasive predator removal are driven by overall reductions in invader abundance or by the suppression of key ecologically impactful life-stages. We argue that this has direct management resource implications and can support effective, targeted, and spatially explicit invasive predator control to maximise conservation gains.

## Methods

### Motivational use case: mink control in Scotland

Large-scale coordinated mink control in Scotland spans over two decades of control efforts split into three phases that vary according to funding, effort structure and scale (Lambin et al., 2019). Phase 1, the Cairngorms Water Vole Conservation Project, started in the lowland in 2004 in the region where our water vole surveys were conducted, and started in the uplands in 2006. Phase 1 was developed to safeguard remnant water vole populations in both lowland and upland regions while relying on volunteers and expanded across c.10,500 km² up until 2009 (Bryce et al., 2011). Following a 19-month with significant reduction in funding that saw limited control and data collection, Phase 2, the Scottish Mink Initiative, was launched in 2011 and lasted until the beginning of 2016 across 29,000 km², approximately a third of mainland Scotland (Lambin et al., 2019). Another two-year funding gap led to a renewed decreases in control and no data collection before Phase 3, the Scottish Invasive Species Initiative (2018 -present) was instated. Phase 3 operates across the same area of Phase 2 using a deliberately thinner monitoring network that focuses on targeting known and newly identified mink breeding sites while gradually expanding into previously uncontrolled areas.

### Data collection

In order to assess whether predator residence status differentially impacts native prey, we use individual locations of mink captured and removed during long-term suppression efforts in Scotland (Lambin et al., 2019) to explore the distance-dependent effect of resident and transient mink on two years of water vole snapshot occupancy.

### Water vole data

Water vole presence-absence data were collected in 2016 and 2023 (Survey 1 and 2 respectively from hereon) within a ∼3,000 km^2^ region within the 29,000 km^2^ mink control area that harboured remnant populations that were steadily declining before mink control started (Strachan et al., 2000; Telfer et al., 2001). Occupancy surveys were conducted along 600 m stretches of waterway sections (sites from hereon) during the water vole breeding season prior to juvenile dispersal (April–June). Presence of water voles was determined by the detection, or not, of latrines (used for marking territories) or freshly used burrows, which have high detectability (Sutherland et al., 2014). Sites were determined *in situ* as occupied (1) if signs were detected or empty (0) otherwise.

In total, 424 sites were surveyed (300 in Survey 1, and 124 in Survey 2; Figure 1). Stratified random sampling was used to select sites in both surveys, maintaining a minimum between- site distance of 200m to ensure spatial independence based on male water vole home ranges (Moorhouse & Macdonald 2005). Only 23.3% (29/124) of sites surveyed in the second survey overlapped with sites from the Survey 1.

**Figure 1.**
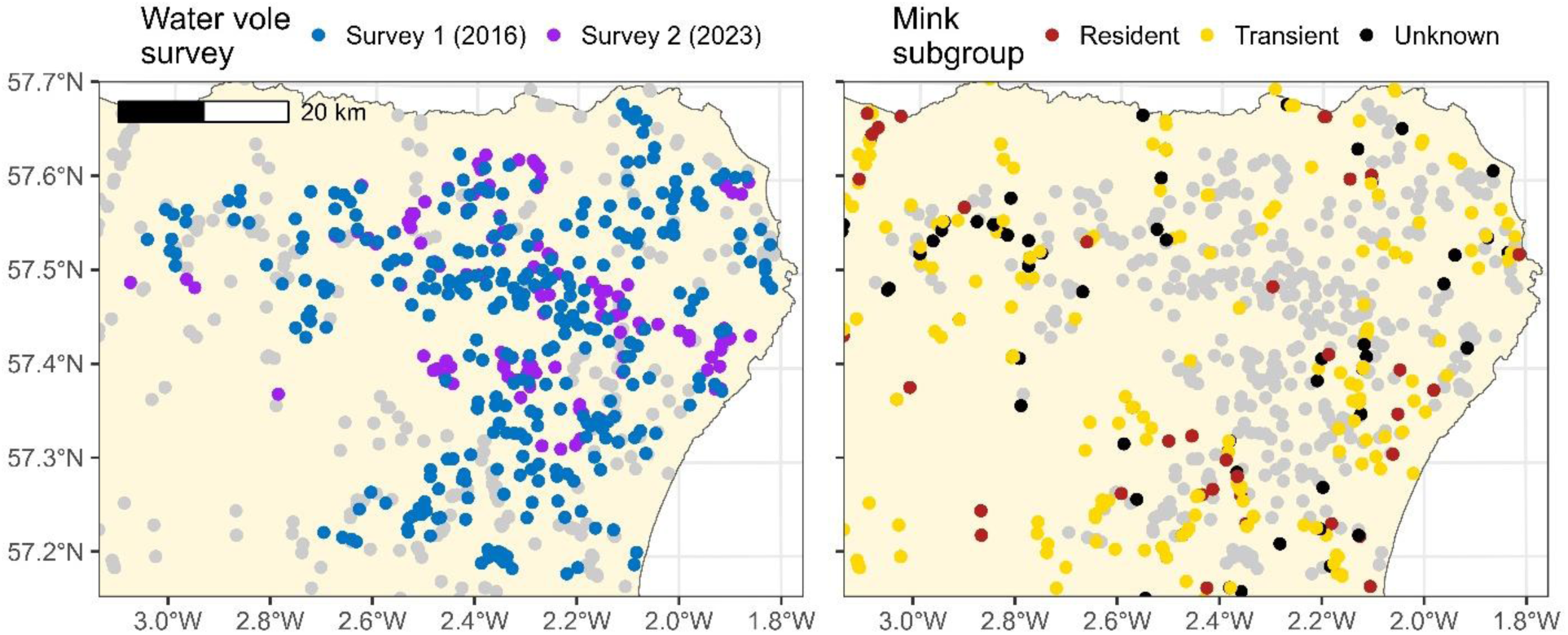
Map showing the distribution of water vole sites (left panel) surveyed during Survey 1 (2016; blue) and Survey 2 (2023; purple) as well as the distribution of mink captures (right panel) of different residence subgroups (resident in red, transient in yellow; unknown in black) within the water vole survey region. In the left hand plot, grey points represent mink captures and in the right hand plot, grey points represent water vole survey sites.

**Figure 2.**
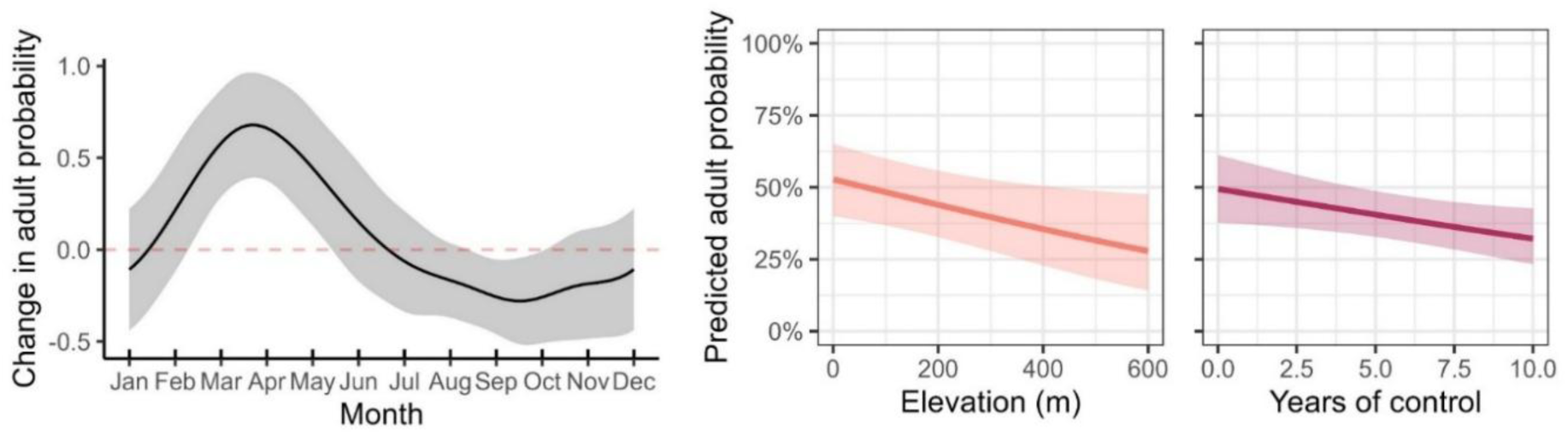
Posterior estimates from generalised additive mink life-stage sub-model results showing the cyclic monthly change in adult capture probability across the months (left panel) as well as the relationship between capturing an adult mink and elevation (pink; right panel) and years of control (dark red; right panel).

Survey 1 was conducted shortly after the end of Phase 2 when regional mink relative density had been reduced from a capture per unit effort of c. 0.27 to less than 0.05 (Oliver et al., 2016). The study area was first divided into 130 hexagonal cells (21.6 km² each, excluding coastal cells with ≤50% of land). Hexagons were then classified into four strata depending on above or below median values of two covariates: waterway density (proxy for mink habitat quality) and connectivity to mink captures following Melero et al. (2015): a) high waterway density and low connectivity, b) high waterway density and high connectivity, c) low waterway density and high connectivity and d) low waterway density and low connectivity. Within each stratum, 60% of the hexagons were selected using randomly selected survey start points ensuring all sites were >200m apart.

Survey 2 occurred five years after Phase 3 of mink control started, during which mink reinvasion, trapping in previously uncontrolled regions and initial deployment in of remotely operated traps in 2023 increased regional capture rates (Lambin et al., 2019). Since water voles were expected to only occur within dispersal/recolonization distance of remnant populations, we defined a sampling frame area of suitable *occupiable* habitat that would minimise the chance of systematic zeros while overlapping with mink captures. To do, we excluded waterways beyond the mean dispersal distance (2.1 km) from any site occupied in at least one of eight previous surveys conducted between 1996 and 2016, i.e. Survey 1 (Lambin et al., 1996; intermediate surveys in 1998, 2002, 2003, 2004, 2005 and 2014).

Sampling units for Survey 2 were river subcatchments (median area 38.8 km^2^; range [4.5 - 116.7]), instead of hexagons. Stratified random sampling was used to using the same four-level stratification, based on waterway density and mean historical mink CPUE, the former proxy for mink connectivity. Subcatchments were randomly selected within each stratum, and survey starting points were randomly allocated to cover ≥30% of each subcatchment’s waterway network.

### Mink capture data

Mink removal during the three control phases followed a sequential approach whereby floating rafts with clay pads were placed in suitable mink habitat to detect mink via footprints (Reynolds et al., 2004). Upon detection of signs of mink presence, a live trap was set and checked daily and removed after a mink was either captured or, typically, after 5 days without mink capture. Some rafts had continuously operational traps when volunteer availability allowed or when the trap was remotely monitored. We use data from 2,025 mink captured across the whole control area from 2008, when Phase 1 trapping network was established, to 2023, the year of water vole Survey 2 (Figure 1). Capture location and date was recorded for every mink captured.

Upon capture, mink were humanely dispatched by staff or trained volunteers and their life-stage was recorded *in situ* after capture by staff members or volunteers if enough determinant signs were observed. Life-stage, defined as juvenile or adult, was determined based on signs of breeding (e.g. raised nipples, mating nape scruff marks, enlarged testes, ossified baculum or penile bone) and measured body length and weight based on staff expertise. In addition to *in situ* observations, the canine teeth of 849 mink captured before 2016 were extracted and aged using tooth cementum annuli analyses performed by Matson Laboratory LLC (MT, USA) using an assumed birthday of April 1^st^. Since mink become sexually mature by the end of their first year (8-12 months), we define their life-stage as adult if they were captured on or after the cut-off of 10 months after April 1^st^ (i.e. February 1^st^). Thus, all other mink captured within 10 months after their assumed birth were defined as juveniles.

### Model outline

In order to investigate how proximity to resident and transient mink influences water voles site-occupancy, water vole patch-occupancy *z_i_* (*z_i_* = 1 if present, *z_i_* = 0 otherwise) in every patch (*i* = 1, 2, 3, …, *N*) was modelled as a Bernoulli random variable:

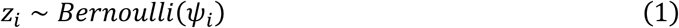

Where *ψ_i_* is the patch-specific occupancy probability modelled as a logit-linear function of site-level covariates:

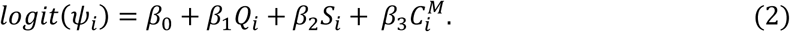

Here *β*_0_ is the intercept, *β*_1_, *β*_2_ and *β*_3_ are the slope coefficients to be estimated for the effects of site-level habitat quality (*Q_i_*), survey (binary; *S_i_* = 1 for sites surveyed during *Survey 2*, *S_i_* = 0 otherwise) and connectivity to mink 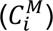, respectively. This formulation means Survey 1 is the reference level and *β*_2_ is the difference in occupancy probability between 2016 and 2023 (Survey 2). Habitat quality is a composite index calculated using microhabitat variables measured *in situ* for both surveys, including variables such as percentage vegetation cover or riverbank structure. The resulting score was calculated using the values from a study by Telfer et al. (2001), which estimated probability of occurrence of water voles in the study system using the same measured covariates. The details of microhabitat measurement and the equation for the composite habitat quality index are in Appendix S1.

As a starting point, we use the framework developed by Chandler & Hepinstall-Cymerman (2016) and reviewed by Yeiser et al. (2021) to calculate connectivity to mink with the assumption that the impacts of individual mink on water vole occupancy decrease with distance between water vole site and mink capture location. Hence, connectivity to mink was defined as the sum of the distance-dependent contributions of all individual captured mink, using a half-normal decay kernel:

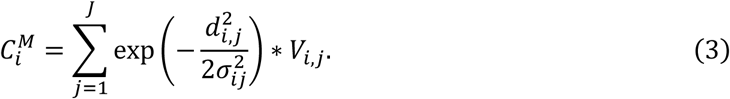

Here, *σ_ij_* is the pairwise scale parameter of the half-normal decay function, determining how rapidly the contribution of mink *j* (*j* = 1, 2, 3, …, *J*) to connectivity 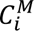 to site *i* decays with squared distance 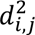 between water vole site *i* and capture location of mink *j*. The larger the *σ_ij_* value is in equation 3, the smaller rate at which mink impacts decay with distance, and thus the broader the range over which individual mink contribute to change in water vole site-occupancy.

To avoid the contribution of mink captured after the surveys were conducted, mink were classified as ‘valid mink’, through the term *V_i_*_,*j*_, a binary indicator that takes value 1 when mink *j* has been captured before the survey of site *i* took place and 0 otherwise.

To be able to incorporate how intraspecific variation in mink influences the spatial scale of connectivity (*σ_ij_*), we extend previous scale-estimating models (Chandler & Hepinstall-Cymerman, 2016) by modelling connectivity decay rate *σ_ij_* as a log-linear function of water vole site-specific and individual mink covariates such that:

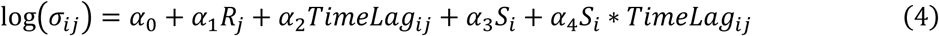

Here *α*_0_ is the baseline log-decay rate for transient individuals and *α*_1_ is the changes in log-decay rates as a function of an individual mink being resident (*R_j_* = 1, 0 otherwise). We also included an effect (*α*_2_) of time lag *TimeLag_ij_* that represents the years between site *i* was surveyed and mink *j* was captured (range: 0-15 for Survey 2 and 0-8 for Survey 1) to account for an expected decrease in effect over time. The term *α*_3_ represents the change in baseline pairwise decay rate when site *i* was surveyed in Survey 2 (*S_i_* = 1) and *α*_4_ is an interaction term that represents the change in time lag effect on Survey 2 in respect to the effect on Survey 1.

### Mink life-stage sub-model

Residence mink status largely depends on the life-stage of individuals to be old enough to have dispersed and established a territory. Gaps in the recording of the life stage of captured of mink (53% missing rate; 1,074/2,025) meant that mink could have been resident unbeknown to us, affecting the estimation of resident-specific scale and magnitude if assumed transient. To circumvent this limitation, we developed and integrated a sub-model that introduces Bernoulli latent mink life-stage (*A_j_* = 1 if adult, *A_j_* = 0 if juvenile) describing the probability of mink *j* being an adult *π_j_* using information from mink with known life-stage to obtain posterior predictions for individuals with unknown life stage and determine their likely resident status. Importantly, we jointly analyse water vole occupancy data and mink-life stage within the same model rather than a two-stage approach to propagate uncertainty around predicted mink life stages.

Once mink with missing life-stage data is known or predicted within the sub-model, the residency status for all mink can be derived if they fulfil at least one of a set of pre-defined conditions. Any adult (*A_j_* = 1) female (regardless of capture date) or any adult individual (male or unknown sex) captured outside the male rutting period (March-April) are classified as resident (*R_j_* = 1). We also consider as resident any juvenile or unknown life-stage caught between November-June, assuming individuals will have completed their dispersal and become resident three months after dispersal events finish in August and before end of weaning in June. Any individuals that do not fulfil any of these conditions these were considered transient (*R_j_* = 0).

The probability *π_j_* of being an adult was modelled as a logit-linear function of covariates that might influence captured mink life stage including capture *Mont*ℎ*_j_* as a cyclic spline to account for fluctuation based on seasonal patterns; elevation (*Elevation_j_*) as a proxy for habitat quality/productivity, as resident individuals are expected to establish in productive lowland (Melero et al., 2018); and control year (*CtrlY_j_*), since the breeding (adult) population was shown to decrease over time (Oliver et al., 2016) such that:

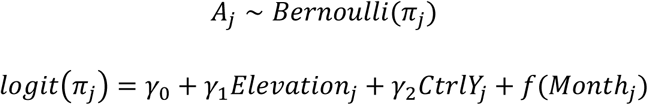

Where the smoother function *f*(*Mont*ℎ*_j_*) represents a cubic cyclic spline modelled as the linear combination of *K (K* = 2 + *l* where *l* is the number of equally spaced knots) spline bases over the range of values within *Mont*ℎ (i.e. 1 to 12) such that:

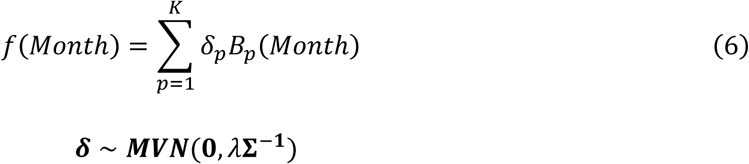

Where ***δ*** is a vector of regression coefficients, one for each of the 1, …, *K* basis splines to be estimated as a multivariate normal distribution with mean 0 and precision matrix equal to the product of smoothing parameter *λ* and inverse variance-covariance matrix **Σ**^−**1**^. Our goal was to estimate settled status-specific connectivity, and its effects on vole occupancy, using known settled status and the posterior predictions from the GAM life-stage sub-model. To ease the estimation process we used the *jagam()* function within the ***mgcv*** (Wood, 2017) to fit the mink life-stage sub-model using those individuals with known life-stage and extract and initial values of ***δ*** and *λ* as well as obtaining the coefficients for **Σ** and value of K (8 here; for more details *see* Wood, 2016).

### Model fitting

The model was fit in a Bayesian framework using Markov chain Monte Carlo (MCMC) algorithm in R v4.4.3 ( R Core Team, 2024) using the ***nimble*** package (v1.3.0;de Valpine et al., 2024) and using the *dbinom_vector()* distribution within the ***nimbleSCR*** package (v0.2.1; Bischof et al., 2022) for the mink life-stage sub-model to speed up computation. To account for correlation between posterior distributions of the kernel parameters, a block random walk MCMC sampler was used for the connectivity distance kernel parameters (Table 1). Vaguely informative normal priors *N*(0, 2.5) were used for the regression coefficients in both the water vole occupancy (Eq. 2) and mink life stage predictors (Eq. 5; Table 1). Three MCMC chains were run with for 100,000 iterations with 60,000 of burn-in and a thinning rate of 2, leading to 15,000 posterior samples per chain. Parameter estimates are reported as posterior means 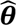 alongside 90% Credible intervals (90% *CrI*) in brackets: 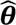 [*LCrI* − *UCrI*]. Convergence was assessed by visual inspection of mixing in trace plots and a potential scale reduction factor *R̂* < 1.05 (Gelman et al., 1995).

**Table 1.**
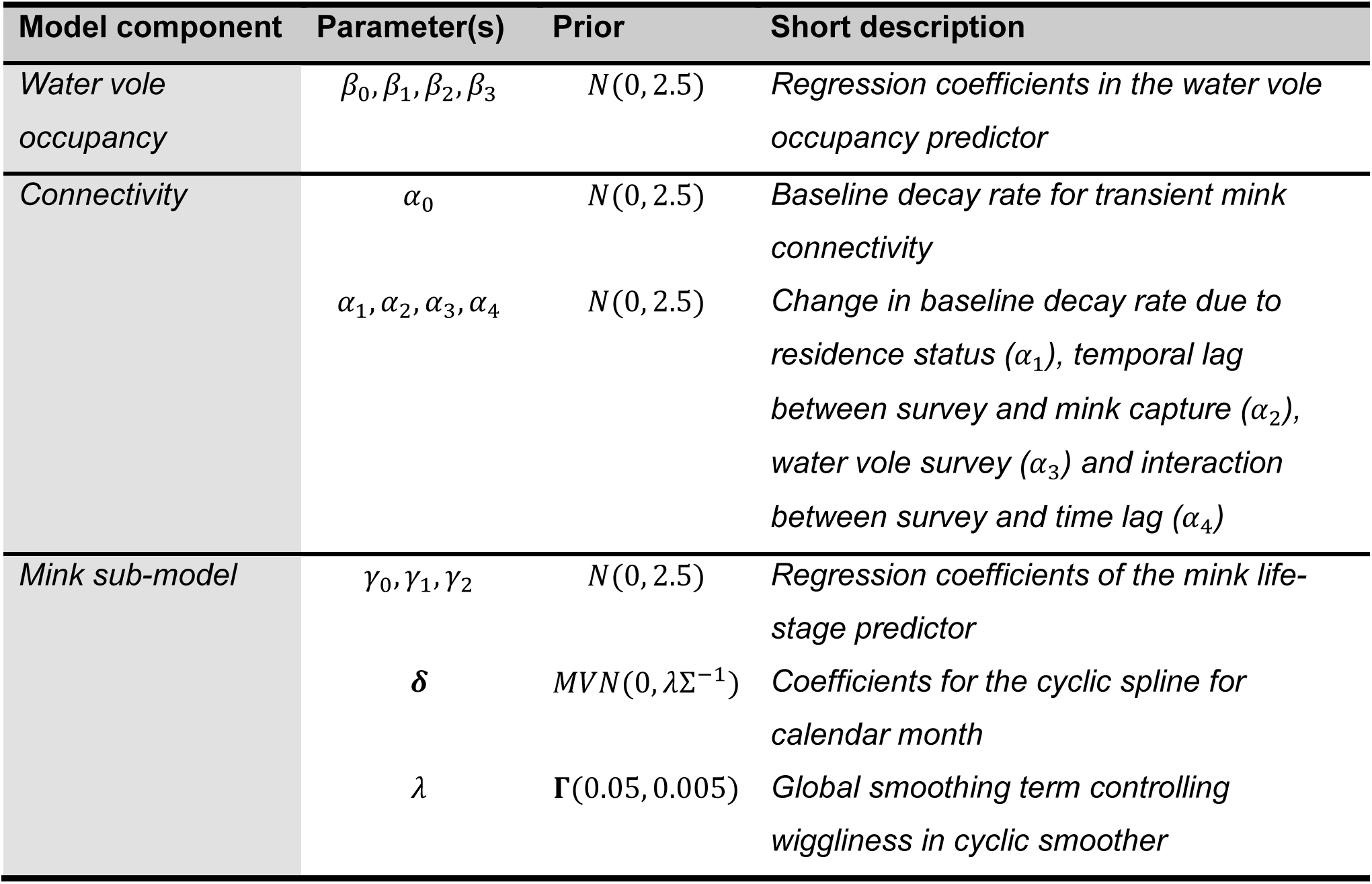
Parameter description and priors for the regression coefficients in the water vole occupancy (β) and mink connectivity scale (α) linear predictors as well as for the mink life-stage sub-model linear (γ) and smoother (δ, λ) regression coefficients

| Model component | Parameter(s) | Prior | Short description |
| --- | --- | --- | --- |
| Water vole occupancy | $\beta_0, \beta_1, \beta_2, \beta_3$ | $N(0, 2.5)$ | Regression coefficients in the water vole occupancy predictor |
| Connectivity | $\alpha_0$ | $N(0, 2.5)$ | Baseline decay rate for transient mink connectivity |
| | $\alpha_1, \alpha_2, \alpha_3, \alpha_4$ | $N(0, 2.5)$ | Change in baseline decay rate due to residence status ($\alpha_1$), temporal lag between survey and mink capture ($\alpha_2$), water vole survey ($\alpha_3$) and interaction between survey and time lag ($\alpha_4$) |
| Mink sub-model | $\gamma_0, \gamma_1, \gamma_2$ | $N(0, 2.5)$ | Regression coefficients of the mink life-stage predictor |
| | $\delta$ | $MVN(0, \lambda \Sigma^{-1})$ | Coefficients for the cyclic spline for calendar month |
| | $\lambda$ | $\Gamma(0.05, 0.005)$ | Global smoothing term controlling wiggleness in cyclic smoother |

## Results

### Mink life-stage sub-model

The baseline probability that a captured mink was adult was 55.8% [49.9-61.4] (*γ*_0_ = 0.234 [−0.005 − 0.464]) and decreased at similar rates with increasing elevation (*γ*_1_ = −0.08 [ −0.12 − −0.03]) and years of control (*γ*_2_ = −0.06 [−0.12 − 0.01]).

The results of the cyclic smooth function within the life-stage sub-model revealed that the baseline probability (*π_j_*) of capturing adult mink varied monthly in a cyclic way (see smoother estimates ***δ*** in the Appendix Table S1). The highest probability of capturing an adult mink was during peak breeding season in and March (71.9 % [59.7 – 81.6]) and April (72.9 % [60.8 – 82.4]) and the lowest after the weaning phase is over and juveniles start dispersing between July (54.3 % [44.0 – 67.9]) and October (49.5% [36.8 – 61.4]).

### Water vole model results

In the absence of effect of proximity to mink captures, the baseline water vole occupancy for Survey 1, under hypothetical zero connectivity effect and average habitat quality, was 77.6% [ 67.9– 85.0] (*β*_0_ = 1.26 [0.75 − 1.74]) and was positively influenced by site-level habitat quality (*β*_1_ = 0.82 [0.63 − 1.01]). Baseline occupancy in Survey 2 was lower (*β*_2_ = −1.54[−2.16 − −0.94], suggesting that in the absence of mink (zero connectivity), baseline occupancy was lower in the second survey (42.4% [28.3 – 57.5]). Connectivity to mink capture locations negatively affected water vole occupancy *β*_3_ = −0.85 [−1.11 − −0.61], reducing water vole occupancy odds by 57.2% [45.7 – 67.0] for every unit increase.

There was strong support for the effect of proximity to mink capture locations to water vole occupancy decreasing slower with increasing distance when captured mink were resident rather than transient (*α*_1_ = 2.34 [ 1.71 − 2.97]; *P*(*α*_1_ > 0) = 100%). Additionally, the effects of both mink subgroups much decayed faster with distance for Survey 2 (*α*_3_ = −2.96 [ −4.44 − −0.94]; *P*(*α*_3_ < 0) = 100%). As expected, the scale of influence decreased for every increasing year between survey and capture year (i.e. temporal lag; *α*_2_ = −0.05 [−0.13 − 0.003]), however there was statistical evidence of an interaction, suggesting that the spatial scale of influence decreased even faster with time lag when sites were surveyed in Survey 1 (*α*_4_ = −1.78[−3.96 − 0.01]; *P*(*α*_4_ < 0) = 90.2%).

For Survey 1, assuming captures occurred in the same year of survey (i.e. time lag was 0) ; the effect of transient mink halved when captures occurred 0.95 [0.31 − 2.74] km from water vole sites, contributing less than 5% by the 1.98 [0.64-5.7] km mark (σ_0_ = −0.21[−1.33 − 0.84]), which suggest their impacts were mostly restricted to the span of water vole sites. Resident mink contribution to connectivity decayed slower, halving at 9.89 [ 5.28 − 22.6] km away from water vole sites and reaching <5% connectivity by the 20.6 [10.9 − 46.9] mark (Figure 4). In contrast, in Survey 2 the contribution of resident and transient mink halved at the 0.29km [0.04 – 1.47] and 0.03 [0.005 – 0.15] km marks (Figure 4), suggesting the distance-dependent effect of mink disappears. This suggests that any influence of mink beyond the immediate survey site was negligible in the second survey (Figure 4). The lack of connectivity effect for Survey 2 means that water vole occupancy was likely driven by habitat quality rather than mink impacts. This in turn explains the lower baseline occupancy for Survey 2 (Figure 3) since the realised negative effect connectivity is negligible compared to that for Survey 1 (Figure 5).

**Figure 3.**
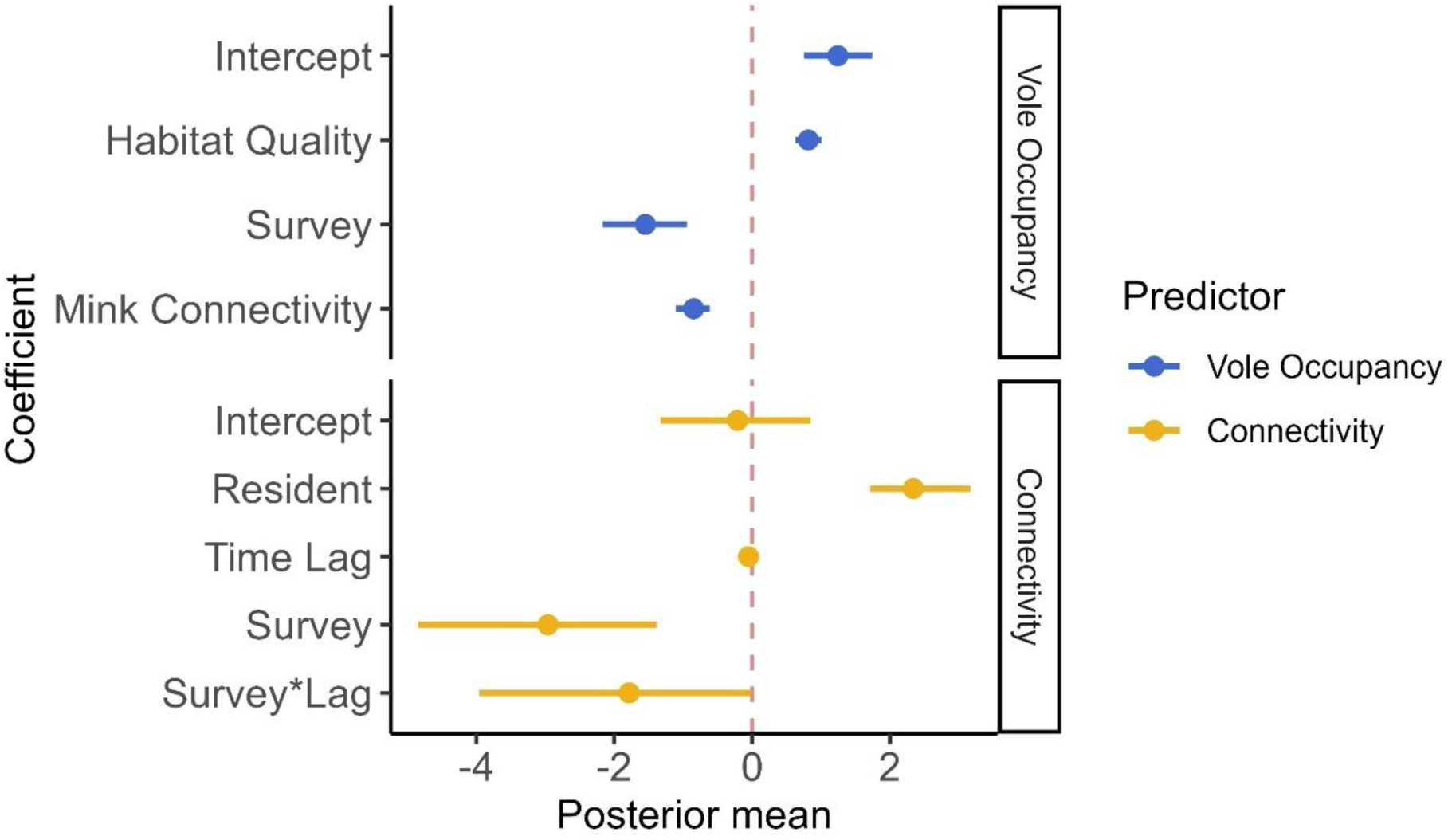
Posterior distribution coefficient regression parameters for the water vole logit-linear predictor (β; blue; top half) and for the vole-mink connectivity scale parameter log-linear predictor (σ; yellow; bottom half). Estimates shown with mean posterior estimate (point) and 90% Credible intervals. Dashed red line at zero to represent overlap of posteriors with zero

**Figure 4.**
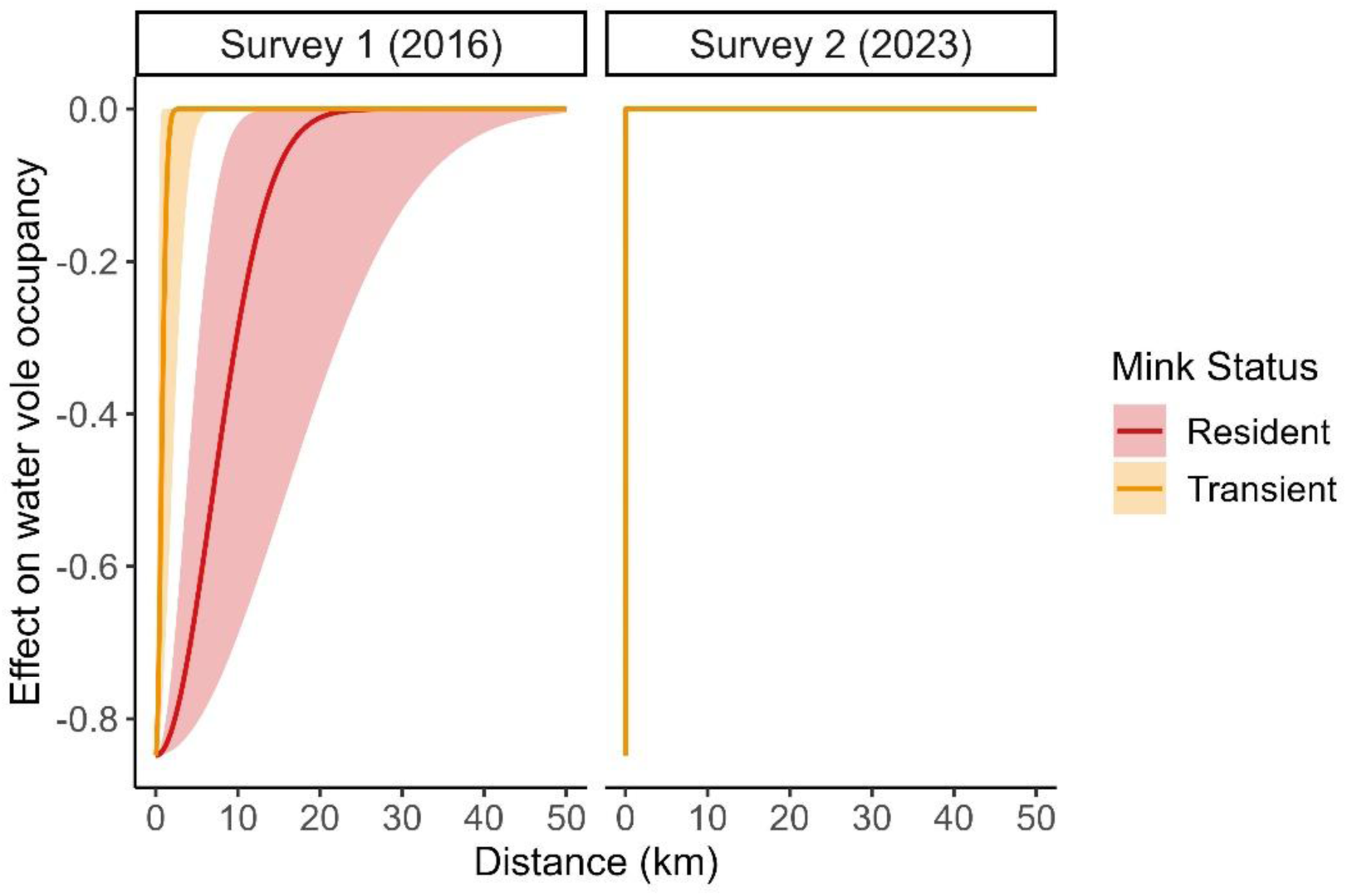
Distance-dependent effect of resident (red) and transient (orange) mink captures on water vole occupancy probability for Survey 1 (2016; left panel) and Survey 2 (2023; right). For Survey 1 the effect of resident mink spans c.10 times further than transient mink. For Survey 2, there is ultimately no spatial signal of effect, leading to the distance-dependent effect to reach zero below the 0.5km mark.

**Figure 5.**
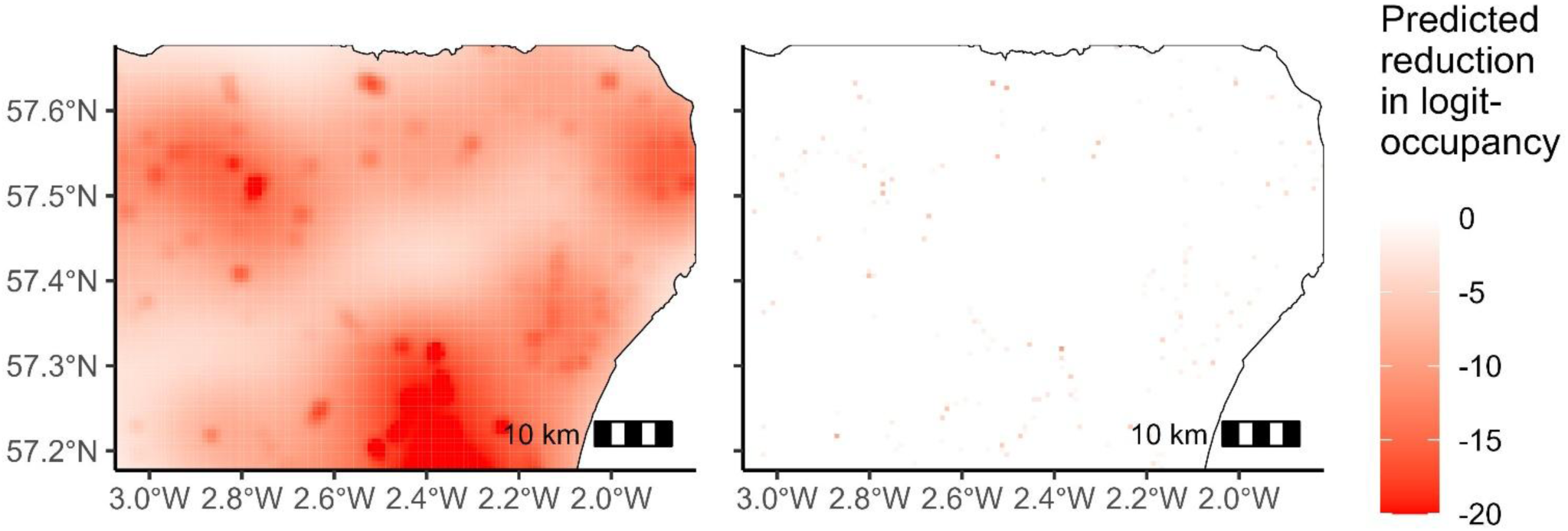
Map of the spatially explicit reduction in water vole occupancy probability (in the logit scale) caused by proximity to resident and transient mink captures for Survey 1 (2016; left panel) and Survey 2 (2023; right panel) across the study area accounting for the effects of all covariates within the connectivity model

## Discussion

Intraspecific heterogeneity in invasive predator impacts is increasingly recognised as an important driver of ecological impacts to be considered when evaluating control strategies (Chinn et al., 2023; Dalal et al., 2021; McCard et al., 2024; Moseby et al., 2015). The impacts of resident mink spanned distances ten times larger than transient mink, providing the first empirical evidence of this previously assumed difference between resident and transient individuals. As control progressed, the distribution of adult resident mink around water vole sites changed, decreasing the spatial scale of influence of mink on water vole occupancy in the second survey. Based on our findings, management actions should evaluate which problem individuals are disproportionally harmful to native populations of conservation concern to design spatially explicit strategies that target their removal.

Our results extend previous evidence gathered from other managed native and invasive predator species that certain individuals or subgroups cause disproportionate ecological impacts. Similar patterns have seen in other invasive mammalian predators like cats (Moseby et al., 2015) and stoats (García-Díaz et al., 2021) as well as in invasive invertebrate predators (European green crabs *Carcinus maenas;* (Kattler et al., 2023) and in native predator subgroups such as dominant coyotes (*Canis latrans*) predating on domestic sheep (Jaeger, 2004). Which introduced predator subgroups can be considered ‘problem individuals’ are likely to be prey-specific, depending on prey size, vulnerability, life-history traits and behavioural adaptability (Barros et al., 2016; Binny et al., 2021). Additionally, landscape contexts such as habitat availability or fragmentation, will influence the ability of prey to respond to predation and removal efforts (Morgan et al., 2022; Rushton et al., 2000). Estimating how predator space use shapes native ecological response to removal may require data too costly to divert from conservation goals. Therefore, we demonstrated how predator data gathered at no additional cost during removal efforts can allow testing of ecological assumptions while providing impact-based management evaluations.

### Residency status influences invasive predator ecological impacts

To date, few studies have explicitly estimated the scale of distance-dependent predator effects on native prey. Previous approaches include modelling how prey abundance changed with distance to invasive lionfish or native goshawks, respectively, as a covariate (Benkwitt, 2016; Mönkkönen et al., 2007). While informative, these approaches do not provide estimates with direct ecological and management interpretation of the spatial scale of predator influence (*σ* in our study), nor do they account for intraspecific impact variations.

Using a novel approach, our results showed that the average spatial scale of resident mink influence (9.89km) on water vole occupancy in Survey 1 was c.10.4 times that of transient mink (0.95 km). This influence extends beyond home range sizes of c.1-6 km estimated for mink (Yamaguchi & Macdonald, 2003; Zabala et al., 2007) and water vole dispersal distances (2.1 km; Sutherland et al., 2014) These findings provide the first empirical evidence confirming expectations that resident mink are the main drivers of water vole declines.

Differences in space use and spatial distribution of predation events between resident and transient individuals, rather than realised ecological impacts, have been reported for native predators (e.g. wolves *Canis lupus* and coyotes; Chamberlain et al., 2021; Hinton et al., 2016). How this translates into ecological impacts is rarely measured due to the difficulty of measuring predation events during dispersal or transience phases, especially from the perspective of prey. Our results provide an interesting insight to this often-unmeasured phenomenon. The limited scale of influence by transient mink suggest that spatially diffuse predation translate into localised impacts within water vole populations, which may not be spatially extensive enough to cause metapopulation-wide declines in occupancy. However, this only provides information on impacts before capture, not across the entire dispersal trajectory, which we expect to be episodic predation impacts across large distances.

Water voles are the flagship species motivating mink control (Lambin et al., 2019), yet mink and other invasive generalist predators have wide diets that vary with sex, size and prey availability. Consequently, within the same predator species which subgroups can be ‘problem individuals’ depends on the prey and socioecological system considered. Female mink are small enough to enter water vole burrows and deplete entire colonies, however larger males predate larger prey species such as ground-nesting birds and rabbits (Craik, 1997; Nordström et al., 2004). In some systems, all mink may be problematic, such as for the critically endangered Hooded grebe *Podiceps gallardoi,* where single incursions from an individual mink killed a significant percentage of the remaining population (Fasola & Roesler, 2018). Indeed, for highly vulnerable prey species in areas of high endemism it will be necessary to remove all individuals since predation impacts may be too large even at low invasive predator densities.

The lack of spatially extensive impacts by transient mink should not be interpreted as that they should not be removed. Logically, removing transient mink before becoming resident prevents new ‘problem individuals to establish in an area, benefiting prey populations. Rather, this suggests that removal of resident mink should be a higher priority. In this case, targeting resident mink will target the breeding population in addition to the more ecologically harmful subgroup, which is key to account for increased fecundity post removal (Melero et al., 2015), the second most important contributor to mink population growth rate after juvenile mortality (Pertoldi et al., 2013). However, individuals responsible for disproportionate impacts may not always be the main contributors to invader population growth rate in other systems (e.g. subadult dingoes in Australia; Henderson et al., 2025). Control efforts should then evaluate whether targeting certain subgroups to benefit one native species will have negative repercussions on supressing the overall invader population, negatively impacting other affected prey species.

### Control context influences scale of predator influence

Control changes the demographic structure of invasive populations, meaning that the realised impact on native prey will also change. Indeed, mink captures after initial depletion phases are dominated by adult males and transient juveniles (Oliver et al. 2016; Melero et al 2017). Our study confirms that adults, the bulk of the resident population, become less likely to be captured as control duration increases. Since water vole extinctions are largely attributed to resident female mink (Strachan et al., 1998), the fact that sustained control depleted resident mink likely reduced the spatially extensive influence of ‘problem individuals’, allowing water vole populations to recover. That would explain the decrease in the spatial scale of influence of mink captures seen in the second water vole survey, in which the mink population is fragmented and composed mostly of transient individuals.

### Utility of the proposed modelling framework

Our modelling framework extended approaches that estimate the spatial scale of landscape effects on local species responses (Chandler & Hepinstall-Cymerman, 2016). Although our method is applied for invasive predator-prey associations, it is broadly applicable to other scale-dependent ecological processes such as competition and reforestation. The distance-dependent term can be incorporated into link function linear predictors of abundance or occupancy, including within state-space frameworks that account for observation error (e.g. N-mixture models; (Madsen & Royle, 2023; Reichert et al., 2017). An appealing feature from our formulation is the flexibility to include species-specific scale-dependent effects, additive or multiplicative, which may be relevant in many systems where native species are simultaneously affected by multiple invasive predators (Doherty et al., 2015).

Another key advantage is that it did not require *a priori* assumptions about the spatial extent of influence, instead estimating scale from the natural expectation that effects decay with increasing distance. Alternative multi-scale or threshold-based methods have been applied to invasive predators (Reichert et al., 2017) and native species (Gallo et al., 2018), but they may underestimate scale of influences or rely on *predefined* scales of influence (Yeiser et al., 2021; Zeller et al., 2012).

Incorporating a sub-model to probabilistically infer missing predator life stage allowed uncertainty to be propagated into estimates of distance-dependent effects on prey occupancy. However, users aiming to replicate this should note that it increases uncertainty in upstream parameters that may risk identifiability, particularly connectivity terms ( *σ_ij_*). Prior-Posterior overlap (PPO) and prior sensitivity checks showed that our parameters were identifiable, but we recommend users validating identifiability when implementing our framework.

For example, it was expected that the effect of mink would vary based on when mink were captured relative to water vole surveys. Under our formulation, we assumed this influenced only the spatial scale of effects, and that when distances between mink and vole sites are zero, mink have the same effect regardless of when captured. This may be unrealistic if ‘ghost of predation past’ effects are mitigated by vole metapopulation (re)colonization and rescue effects. This could be modelled by modifying Eq. 3 so the zero-distance effect depends on mink residence status and temporal lag. However, we found this formulation unfeasible with our data. Preliminary analyses showed identifiability issues, with PPO values above 50% for all, reaching 72%, 89%, and 70% for some connectivity scale and intensity parameters (Appendix Figure S2). We believe this lack of identifiability may reflect correlation between baseline and scale parameters, where multiple combinations yield the same expected water vole occupancy probabilities. This issue may be reduced when data is abundant at multiple distances and temporal lags, allowing clearer separation of effects. Reduced mink abundance at the end of Survey 1 and throughout Survey 2 led to fewer recent captures near vole sites, lowering the effective sample size at zero distances and the ability of the model to estimate all parameters. However, further research is needed to determine whether this issue is structural or data-specific, and whether other formulations may circumvent it.

### Management implications

Our work supports the development of targeted spatially explicit impact-based management for effective resource allocation in control efforts. Managers can use spatially explicit predictions of invasive species impacts to create buffers around native species sites where impacts are expected to be greatest. This can be particularly beneficial in reintroduction campaigns using man-made ‘safe-havens’ where predation ‘beyond the fence’ threatens released population, where managers may need to create a safety buffer around reintroduction areas to increase reintroduction success (Smith et al., 2023). For our case study, to minimise mink impacts on water voles, managers should prioritise removing resident mink within c.10km of water vole sites, but ideally up to 20km. This will be especially relevant in initial phases of removal, where targeted removal of resident mink at high initial densities will maximise reduction of ecological impacts while depleting invader abundance. Efforts then can be concentrated in habitats attractive to resident mink (Melero et al., 2018) or use trapping methods that increase resident mink capture rates, such as gland scent lure or remotely operated devices (Martin, 2022).

As control progresses, managers should re-evaluate resource allocation and the effectiveness of impact-based metrics, especially in long-term conservation efforts where landscape context, and species interactions change through management. Adaptive management strategies that iteratively update evidence as new data is collected can be beneficial in invasive predators and broader conservation management (Blomquist et al., 2010; Walters & Holling, 1990). For territorial predators, home range sizes may increase as control reduces density (Effort et al 2004), potentially increasing the realised impacts on prey as seen for invasive stoats (García-Díaz et al., 2021). Control buffers required to maintain native species recovery will therefore change over time. Removal can also increase the emergence of ‘problem individuals’ when non-problem individuals can establish in newly emptied area or, for example, can rise in social hierarchy (e.g. subdominant coyotes becoming dominant; Jaeger, 2004). In those cases, management should remove all individuals to prevent individuals from becoming problematic. For mink, establishment by transients increases as dispersal distances to suitable habitat decrease following resident mink removal, especially if fecundity rates increase post-removal (Melero et al. 2015; Oliver et al. 2016). Increasing trapping effort during the initial months of post-natal dispersal, when juveniles emerge from natal territories can help limit the rise in establishment rates.

While targeted removal can outperform blanket removal when impacts are driven by a subset of individuals (Lennox et al., 2018; Swan et al., 2017), managers should evaluate whether it is feasible in their system. This strategy requires problem individuals to have traits that can be reliably identified (Swan et al., 2017). In our case, problem mink was linked to life-stages and residence status and similar predictability has been identified in other species such as juvenile brown bears (Elfström et al., 2014) or large male feral cats (Moseby et al., 2015). However, problem individuals may arise from behavioural specialisation that cannot be linked to consistent traits over time, making them challenging to detect.

Lastly, managers should consider which native species are appropriate for evaluating abundance-impact relationships. For generalist invasive predators, targeting specialist predators may benefit only one of multiple affected species. Additionally, in multi-predator invaded systems removal of one predator may reduce interspecific competition, inadvertently undermining conservation efforts if predation pressure by non-target invasive predators increases through ‘mesopredator release’ (Doherty et al., 2015). We advocate that targeted removal can be a powerful strategy to maximise conservation gains, but managers must carefully consider whether it is applicable in their system. In our study, invasive mink are the only invasive predator threatening water vole persistence. In systems with multiple invasive predators or where habitat for native prey is too fragmented (Morgan et al., 2022), an overinvestment in targeted removal may detract from the fundamental conservation objectives to reduce invasive predators abundance and allow native species to recover.

## Acknowledgements

A.B.B. has received funding under the NERC Scottish Universities Partnership for Environmental Research (SUPER) Doctoral Training Partnership (DTP) (Grant reference number NE/S007342/1) and NatureScot. X.L. was in receipt of a Leverhulme fellowship RF-2024-363. We thank Dr. Ewan K. McHennry and Dr. William H. Morgan for their data collection efforts for the 2016 data. We thank all staff and volunteers in the Scottish Invasive Species Initiative and associated partner organizations, including but not limited to Callum Sinclair, Jane Hamilton and Chris Horril (deceased). We are grateful to Dr. Jason Mathiopoulos and Dr. Sandra Telfer for providing valuable comments of this manuscript.

## Author contributions

A.B.B: Conceptualization, Data curation, formal analyses, Investigation, methodology, project administration, validation, visualization, writing – Original draft, writing – review & editing; C.S.: Conceptualization, Funding acquisition, Investigation, Supervision, Writing – Review & editing; X.L.: Conceptualization, Funding acquisition, Investigation, Project administration, Supervision, Writing – Review & editing

All authors gave final approval for publication and agreed to be held accountable for the work performed herein.

## Artificial intelligence use declaration

Artificial intelligence was used for grammatical proofreading of finalised draft only (Grammarly AI). We declare no use of AI in the conceptualization, analyses, visualization and original writing.

## Data availability statement

Data and code is available in an online repository within the Open Science Framework: https://osf.io/y9j4e/overview?view_only=0f00612537cc45a4a6ab530c0b1d095b

