## Appendix Table S1 for "One scale does not fit all: invasive predator identity determines the impact on native prey"

**Supplementary material**

### Habitat quality index

To derive habitat quality at the site level we used the regression coefficient values that represent the change in occupancy probability based on given environmental covariates derived from an occupancy study conducted by Telfer et al. (2001) in our same study area before mink control started (see equation below). Within a 600m site, the covariates were measured at the 100m, 300m and 500m marks representing the middle point of 200m sections that approximate the home range of male water voles. The variables were the following riverbank vegetation cover (%;*Veg*) in a 2m^2^ quadrat; tree cover (%;*Tree*); *Juncus effusus* (%; *Ju*) and *Filipendula ulmaria* (%;*Fu*) cover ; riverbed substrate type (0-mud, 1-gravel, 2-rock; *Sub*); and bank penetrability divided into 3 classes (Pen1-soft, Pen2-medium, Pen3-hard) measured by inserting a metal pole of 2.5cm diameter into the riverbank. Habitat quality was calculated for each section using the equation below and average site quality was the mean across the three sections’ scores.

$$HQ= 0.069Ju -0.095Fu-0.029Tree+7.39Pen1+7.20Pen2+4.92Pen3+0.019Veg -2.53*Sub$$

### Smooth parameter estimates

*Table S1* Posterior mean and standard error estiamtes for the cyclic cubic splines ($\delta$) and global penalty term ($\lambda$) within smoothing term in the mink life-stage sub-model that describes how mink adult probability changes with month

| **Parameters** | **Notation** | **Mean** | **Standard error** |
| --- | --- | --- | --- |
| *Basis spline coefficients (8)* | $\boldsymbol{\delta}$ | $\boldsymbol{\delta=}$( $-0.58,-0.34,-0.06,0.11,$ $-0.19,-0.25,0.38, 0.55$) | $\boldsymbol{SE}\left( \boldsymbol{\delta} \right)\boldsymbol{=(}0.18,0.13, 0.15, 0.17,$  $0.16, 0.14, 0.15, 0.13\boldsymbol{)}$ |
| *Global penalty smoother* | $\lambda$ | $\hat{\lambda}=$34.7 | $SE\left( \lambda\right)=27.4$ |

### Intensity lag identifiability issues

It is expected that the effect of mink on water vole occupancy would change depending on when mink were captured relative to the water vole survey. Under our formulation, we assume that this effect is within the scale of mink influence, that being how far in space the negative association between mink and vole spans. This assumes that, when distance between mink and vole sites is zero, mink have the same effect on vole occupancy regardless of when these were captured. In truth, this is likely unrealistic. Through colonization-extinction dynamics in vole metapopulations, the intensity of the effect of ‘ghost of predation past’ by mink may be rapidly compensated through (re)colonization and rescue effects. In practice, this temporally varying intensity could be modelled by modifying Equation 3 to include a ‘baseline intensity’ parameter, where the effect of mink at distance zero ($g_{0j}$) is modelled as a logit-linear function of, for example in our system, residence status and temporal lag:

$$C_{i}^{M}=\sum_{j=1}^{J} g_{0j}*exp \left( -\frac{d_{i,j}^{2}}{2\sigma_{ij}^{2}} \right)*V_{i,j}.$$

$$logit(g_{0j})=\alpha_{0}+\alpha_{1}R_{j}+\alpha_{2}TimeLag_{j}$$

Where $\alpha_{0}$ is the baseline effect of transient mink ($R_{j}=0$) captured in the same year as the water vole survey ($TimeLag_{j}=0$), $\alpha_{1}$ is the difference in baseline effect when the mink is resident ($R_{j}=1$) and $\alpha_{2}$ is the change in baseline effect per year in the past between mink capture and vole survey. The rest of parameters are the same as Equation 3 in the main text.

However, this formulation was unfeasible from an estimation point of view. We found in preliminary analyses aiming to model both how the intensity of effect ($\alpha$) and spatial scale of mink capture effects varied with time, that the connectivity parameters become unidentifiable. With prior-posterior overlaps (PPO) of 72%, 89% and 61% for the $\boldsymbol{\alpha}$ parameters, and 70%, 52% and 51% for the $\sigma_{0}$, $\sigma_{3}$ and $\sigma_{4}$ respectively (Figure S1). We believe the main explanation for this lack of identifiability is the structural correlation between baseline and scale parameters, where multiple combination of parameter values could lead to the same expected water vole occupancy probabilities. Whilst this correlation could be mitigated with sufficient data at the multiple distance and temporal bands. Reduced mink abundance at the end of Survey 1 and throughout Survey 2 led to fewer recent captures near vole sites, lowering the effective sample size at zero distances and the ability of the model to estimate all parameters.

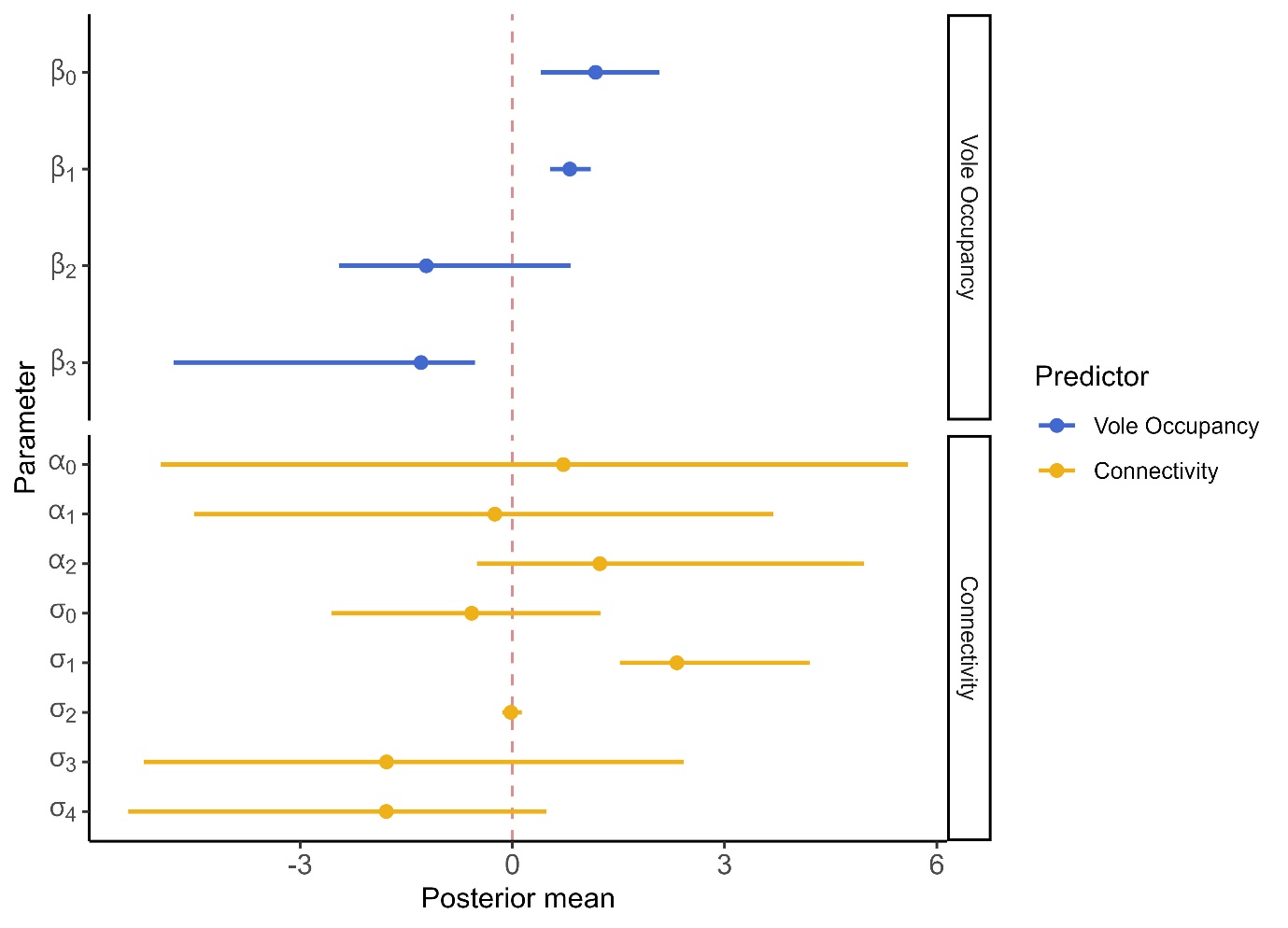

*Figure S1 Posterior mean and 95% Credible Estimates obtained in preliminary analyses of a more complex of the model fitted to estimate distance-dependent effects of transient and resident mink on water vole occupancy. Here we include and model the intensity of effect of mink (* $\alpha$*parameters) as a function of residency status and time lag between mink capture and vole survey.*
